# Evaluation of out-of-distribution detection methods for data shifts in single-cell transcriptomics

**DOI:** 10.1101/2025.01.24.634709

**Authors:** Lauren Theunissen, Thomas Mortier, Yvan Saeys, Willem Waegeman

## Abstract

Automatic cell type annotation methods assign cell type labels to new, unlabeled datasets by leveraging relationships from a reference RNA-seq atlas. However, new datasets may include labels absent from the reference dataset or exhibit feature distributions that diverge from it. These scenarios can significantly affect the reliability of annotation predictions, a factor often overlooked in current automatic annotation methods. The field of out-of-distribution detection (OOD), primarily focused on computer vision, addresses the identification of instances that differ from the training distribution. Implementing OOD methods in the context of novel cell type annotation and data shift detection for single-cell transcriptomics may enhance annotation accuracy and trustworthiness. We evaluate 6 OOD detection methods: LogitNorm, MC dropout, Ensembles, Energy-based OOD, Deep NN and Posterior networks, for their annotation and OOD detection performance in both synthetical and real-life application settings. We show that OOD detection methods are able to accurately detect novel cell types and show their promise to detect severe data shifts on non-integrated datasets. Moreover, integration of the OOD datasets improves annotation performance, but interferes with OOD detection, diminishing novel cell type capabilities of the OOD methods.

## 1 Introduction

The development of automatic cell type annotation tools has been a popular area of research in recent years. Large-scale atlases Regev et al. [2017], Sikkema et al. [2023], Deprez et al. [2020] have significantly enhanced the value of automatic cell type annotation, as they can be used to build pre-trained annotation tools Hrovatin et al. [2024]. End-users can use these pre-trained tools Li et al. [2023], Hou and Ji [2024], Shao et al. [2021], Yang et al. [2023] to assign cell type labels to their own, often smaller and more specific, datasets and leverage the rich information present in the atlases. However, these smaller datasets can contain biological and or technical variation that is not present in the training atlases, due to the presence of new labels, patients, disease states, tissues, the use of different protocols, etc. This often results in a distribution or data shift between the reference data, here the atlas, and the test data, the smaller dataset. Data shifts severely impact the performance and reliability of the tools as generally multi-class classification models – such as cell type annotation tools – assume that training and test datasets are independent and identically distributed (i.i.d.) according to an unknown distribution. In practice, these distribution shifts can lead to a much higher number of incorrectly allocated cell type labels than the end user might anticipate, given the tools’ reported performance Sun et al. [2022a]. Ideally, any annotation tool should be able to accurately detect the presence of novel cell types in the test dataset, but also detect data shifts that severely influence the tool’s annotation performance, so that the data shift can be mitigated with the help of integration techniques Luecken et al. [2022] or studied in more depth.

Out-of-distribution (OOD) detection is an interesting machine learning field that tries to handle the effects of data shifts during classification. OOD detection methods intend to identify and flag samples in the test data that are affected by data shifts, ensuring accurate and reliable classification ingkang Yang et al. [2024]. Numerous OOD detection methods have been developed, particularly in the field of computer vision Yang et al. [2022, 2024], Liu et al. [2023], Cui and Wang [2022], Lu et al. [2024], Salehi et al. [2022]. These methods in essence try to perform uncertainty quantification and threshold their uncertainty estimates to identify and flag aberrant samples, i.e., samples deviating from the training data distribution. In uncertainty quantification literature, the total predictive uncertainty is often divided into two components: aleatoric and epistemic. Aleatoric uncertainty arises due to inherent randomness in the data generating process and cannot be reduced with more training data. Epistemic uncertainty, on the other hand, is uncertainty about the optimal model parameters and can be mitigated by adding more data. Hüllermeier and Waegeman [2021], Charpentier et al. [2020]. In theory, OOD detection methods should base their OOD decisions on epistemic uncertainty only, but this is not always the case.

In the single-cell field, several annotation tools incorporate a reject option to manage data shifts and flag the subsequent hard-to-recognize instances or novel cell types Michielsen et al. [2021], Conde et al. [2022], Fischer et al. [2024], Ma and Pellegrini [2020], Kiselev et al. [2018], Zhang et al. [2019]. These reject options are based on various mathematical concepts, few from the OOD field, and are tested on specific use cases, making it hard to gauge their overall performance. Currently, to the best of our knowledge, only one preprint compares the OOD performance of two OOD-specific (methods specifically designed for OOD detection) and two non-OOD-specific methods, but this evaluation is limited to a synthetic OOD setting with only three excluded cell types Engelmann et al. [2022]. A comprehensive comparison of different OOD methodologies for cell type annotation in single-cell transcriptomics, across various real-life biological settings, is still lacking.

In this paper, we evaluate six established OOD methodologies from the computer vision field for detecting data shifts and novel cell types in single-cell transcriptomics across three datasets. We assess these methodologies based on their in-distribution (ID) annotation performance and their OOD detection capabilities. For OOD detection, we synthetically remove the least common cell type populations during training, as is commonly done in the literature, but also examine various applications with naturally occurring data shifts, such as annotating data from a new patient, tissue or disease or data generated with a different protocol. We also explore how these biological shifts impact cell type annotation. Finally, we investigate the impact of integration on cell type annotation and OOD detection.

## 2 Material and Methods

### 2.1 Formal problem definition

**Cell type annotation** is a multi-class classification problem, where the goal is to predict cell type labels from the label space 𝒴 = *{*1, …, *K}*, which contains *K* classes, based on inputs from some input space 𝒳. The training dataset 𝒟 _train_ = *{*(**x**_1_, *y*_1_), (**x**_2_, *y*_2_), …, (**x**_*N*_, *y*_*N*_)*}* contains *N* data points, sampled i.i.d. from some unknown joint distribution 𝒫 _train_(**x**, *y*) on 𝒳 × 𝒴. Given 𝒫 _train_, we will learn with most methods a classifier that returns probabilities. More formally, this is a classifier *f* : **x** → Δ^*K*^ with 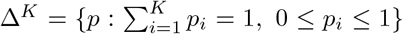. The classifier *f* will be optimized with the help of a loss function ℒ (·).

**Dataset shifts** occur when the training and test joint distributions are different 𝒫_train_(**x**, *y*) ≠ 𝒫_test_(**x**, *y*). Dataset shifts can be categorized into three different categories: covariate shifts, prior probability shifts and concept shifts. Covariate shift, also called population drift, refers to a change in the distribution of the input variable **x**: *P*_train_(*y*|**x**) = *P*_test_(*y*|**x**), but *P*_train_(**x**) ≠ *P*_test_(**x**).A prior probability shift refers to changes in the distribution of the label space *Y*: *P*_train_(**x**|*y*) = *P*_test_(**x**|*y*) but *P*_train_(*y*) ≠*P*_test_(*y*). A concept shift or drift occurs when there is a change between the relationship of 𝒳 and 𝒴. Formally, this can be defined as *P*_train_(*y*|**x**) *P*_test_(*y*|**x**) but *P*_train_(**x**) = *P*_test_(**x**) Moreno-Torres et al. [2012]. In single-cell transcriptomics, covariance shifts can happen due to various biological factors. Occasionally, these shifts are accompanied by a prior probability shift because of new cell type labels in the test data. A concept shift is not naturally present in the setting of cell type annotation for single-cell transcriptomics and will thus not be considered in this evaluation (see Section Datasets).

**Out-of-distribution detection** is a binary classification problem, where the goal is to label an input **x** ∈ 𝒳 as in-distribution (ID) if it belongs to 𝒫 _*train*_ and out-of-distribution (OOD) if it does not. In practice, OOD detection is often implemented with the help of a scoring function *S*(**x**) so that:

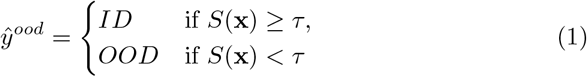

where *τ* is a user-defined threshold and *ŷ*^*ood*^ the OOD prediction. In this paper we evaluate the methods’ ability to both correctly perform ID classification, as well as to correctly perform OOD classification. This setting is also referred to as open-set recognition, open category detection or open set learning in the machine learning literature, but not all authors adhere to this nomenclature ingkang Yang et al. [2024].

### 2.2 OOD Methods

In this paper we evaluate six OOD methods: LogitNorm Wei et al. [2022], MC dropout Gal and Ghahramani [2016], Deep ensembles Lakshminarayanan et al. [2017], Energy-based OOD (EBO) Liu et al. [2020], Deep nearest neighbors (Deep NN) Sun et al. [2022b] and Posterior Networks Charpentier et al. [2020]. We chose these methods based on their performance in the recent out-of-distribution benchmark by Yang et al. (2022) Yang et al. [2022] and their popularity. We categorized these methods into 3 categories based on the mathematical principles they use to detect out-of-distribution samples (softmax-based methods, density-based methods and distance-based methods) and also on their open-worldness during training (see Table 1). LogitNorm, MC dropout and Deep ensembles all use the final softmax output score of the classifier *f* to perform OOD detection. Energy-based OOD and Posterior Networks construct densities, based on the training (input) data, and evaluate the test data w.r.t. these training densities. Deep NN performs distance-based OOD by using the distances between the test and training data to perform OOD detection. Posterior Networks, Energy-based OOD and Deep NN explicitly account for open world observations during training, i.e. they take the possibility of novel, unseen labels into account during their training phase.

**Table 1:**
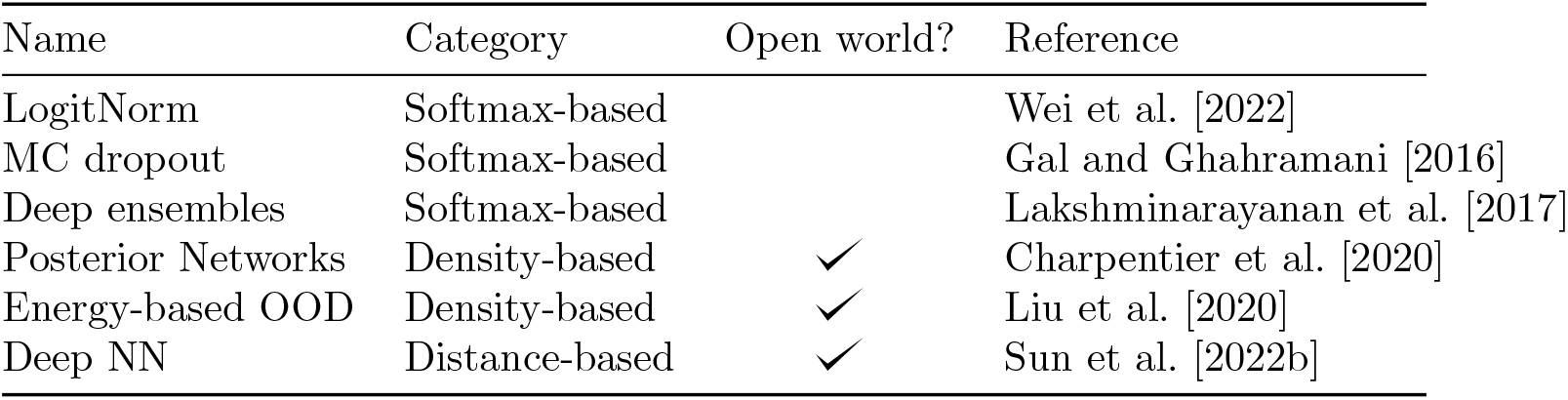
Overview and categorization of the six implemented OOD methods.

| Name | Category | Open world? | Reference |
| --- | --- | --- | --- |
| LogitNorm | Softmax-based |  | Wei et al. [2022] |
| MC dropout | Softmax-based |  | Gal and Ghahramani [2016] |
| Deep ensembles | Softmax-based |  | Lakshminarayanan et al. [2017] |
| Posterior Networks | Density-based | ✓ | Charpentier et al. [2020] |
| Energy-based OOD | Density-based | ✓ | Liu et al. [2020] |
| Deep NN | Distance-based | ✓ | Sun et al. [2022b] |

**LogitNorm** The authors of the LogitNorm method start from the assumption that the model should produce predictions with lower confidence scores for OOD data, compared to ID data Wei et al. [2022]. However, in reality, the confidence scores are often overconfident, i.e., high scores are assigned to all predictions regardless of their correctness. This problem mainly occurs in deep neural networks Guo et al. [2017]. We will use *h*(**x**) to denote the outputs of the penultimate layer of the neural network *f* (**x**). These outputs are commonly referred to as the logits. The authors consider the cross-entropy loss ℒ_*CE*_:

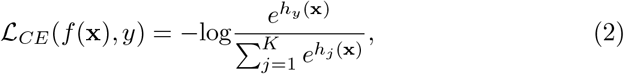

to optimize the NN model *f* and show that the overconfidence of the model is caused by the cross-entropy loss that keeps increasing the magnitude of the logit vectors, even when a sample is already correctly classified. To alleviate this problem the Logit Normalization loss, dubbed the LogitNorm loss ℒ _LogitNorm_, is proposed as an alternative to the cross-entropy loss:

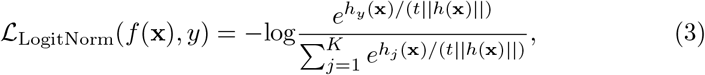

where *t* The logits will modulate the magnitude of the logits *h*(**x**). *h*(**x**) can be decomposed into two components without loss of generality, the euclidean norm ||*h*(**x**)||, which represents the magnitude of the logit vector, and the unit vector 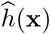, which depicts the direction. The LogitNorm loss ℒ _LogitNorm_ (Eq.3) decouples the influence of the logits’ magnitude from the training process Wei et al. [2022]. The scoring function used for OOD detection is derived from the probabilities returned by the neural networks *f*, which was trained with the logitnorm loss ℒ_LogitNorm_: *S*(**x**) = max_*k*=1,…,*K*_ *f*_*k*_(**x**).

**MC dropout** is a well-known randomization technique that was initially introduced to improve predictive performance, but nowadays it is also commonly used for uncertainty quantification and OOD detection. With the Dropout mechanism, each node of each layer is dropped or excluded from the NN during training with a dropout probability of 0.5. In this way, for each training sample a different thinned network is sampled and trained Srivastava et al. [2014]. Gal et. al (2016) Gal and Ghahramani [2016] show that an NN, with Dropout applied before every weight layer, can be interpreted as a Bayesian approximation of the Deep Gaussian Process model. With MC dropout, *T* stochastic forward passes are made through the dropout network. So *T* thinned networks are sampled and evaluated. The final prediction scoring function, used for OOD detection and cell type annotation, is the softmax of the averaged logits over the *T* forward passes or evaluated networks: 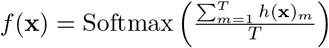 and *S*(**x**) = max_*k*=1,…,*K*_ *f*_*k*_(**x**) Gal and Ghahramani [2016].

**Deep Ensembles** improves model uncertainty estimation by using an ensemble of NN-models, trained with a proper scoring rule^1^such as the cross entropy loss ℒ _*CE*_ (Eq. 2) Lakshminarayanan et al. [2017]. With Deep Ensembles *T* NN-models are randomly initialized, independently trained, and the softmax averages of the *T* logits *h*(**x**) are used for OOD detection and cell type annotation: 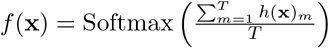 and *S*(**x**) = max *f* (**x**).

**Posterior Networks** explicitly model an (epistemic) uncertainty distribution ***q*** characterized by the family of Dirichlet distributions, over the categorical class distribution **p** = [*p*_1_, …, *p*_*K*_] that is estimated by *f* (**x**) for every **x**. Deep ensembles and MC dropout represent ***q*** implicitly, and can only sample from it and estimate statistics on it. The explicit parametrization of this uncertainty distribution ***q*** over the categorical class distribution ***p*** allows to compute and distinguish between epistemic uncertainty, aleatoric uncertainty and class predictions in one pass. Posterior Networks map **x** to a latent space *Ƶ* with the help of an encoder network *f*_*Encoder*_. They also train a Normalizing flow, parameterized by *ϕ*, to learn flexible density functions *P* (***z*** |*ϕ*) for every class, that are evaluated at the positions of the latent vectors ***z*** ∈ Ƶ. The resulting densities are used for the parametrization of the Dirichlet distribution ***q*** for each data point ***x***. Higher densities correspond to higher confidence in the Dirichlet distribution. When ***x*** corresponds to a low density region, a very high epistemic uncertainty will be predicted by the Dirichlet distribution. Both the encoder network and the normalizing flow are jointly optimized with the help of the uncertainty-aware loss ℒ_UCA_ Charpentier et al. [2020]:

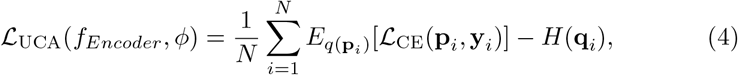

where *N* indicates the number of observations, **y** the one-hot-encoded vector representation of *y* and *H*(**q**), an entropy regularizer that promotes smooth distributions of **q**. The normalized densities for every sample are used to perform OOD detection and cell type annotation. Class probabilities are obtained from the densities by applying Bayes’ rule, while the OOD detection score *S*(**x**) is proportional to the highest class density.

**Energy-based OOD** is a post-hoc method that consists of calculating an energy-score *S*_*E*_ after training and inference based on the logits *h*(**x**) returned by the NN model *f* :

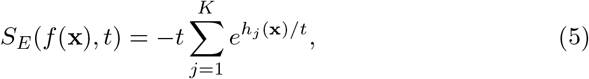

where t is a temperature parameter that rescales the logits *h*(**x**). Samples with higher energy scores will be seen as more likely OOD, thus the negative values of the energy scores will be used for OOD detection Liu et al. [2020].

**Deep Nearest-neighbors** The Deep NN method performs non-parametric density estimation with the help of k-nearest neighbors. The method calculates the Euclidean distance in the space formed by the normalized penultimate layer embedding: *h*_*norm*_(**x**) = *h*(**x**)*/* ∥*h*(**x**) ∥_2_. This is done for every observation of the test data 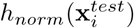 and its *k*-th nearest neighbor **x**_*k*_ in the normalized- penultimate embedded training data 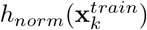. This distance is then used as a score for OOD detection Sun et al. [2022b].

### 2.3 Datasets and model construction

We evaluated the OOD methods across 3 different datasets: the Lung, Immune and COPD dataset [Luecken et al., 2022, Adams et al., 2020]. Information on the preprocessing of the datasets can be found in Appendix 1.1. The Lung dataset consists of three healthy 10X transplant datasets, one Drop-seq transplant dataset and one 10X lung biopsy dataset. The Immune dataset contains peripheral blood mononuclear cell (PBMC) data, sequenced with SMART-seq and 10X, together with Bone marrow data sequenced with 10X. The COPD dataset contains samples from healthy (smoker and non-smoker) patients, patients with idiopathic pulmonary fibrosis (IPF) and patients with chronic obstructive pulmonary disease (COPD), all sequenced with 10X. Inspection of the COPD dataset showed no clear disease (batch) effects between the disease and control data parts, resulting in our believe that the dataset is most likely integrated (Figure 1A.), though this is not clearly specified by the authors in the manuscript or data file [Adams et al., 2020]. Each OOD method was implemented with multiple neural network (NN) architectures for *f*. The results of the best overall performing networks are reported, for more information on model construction we refer the reader to Appendix 1.2.

**Figure 1:**
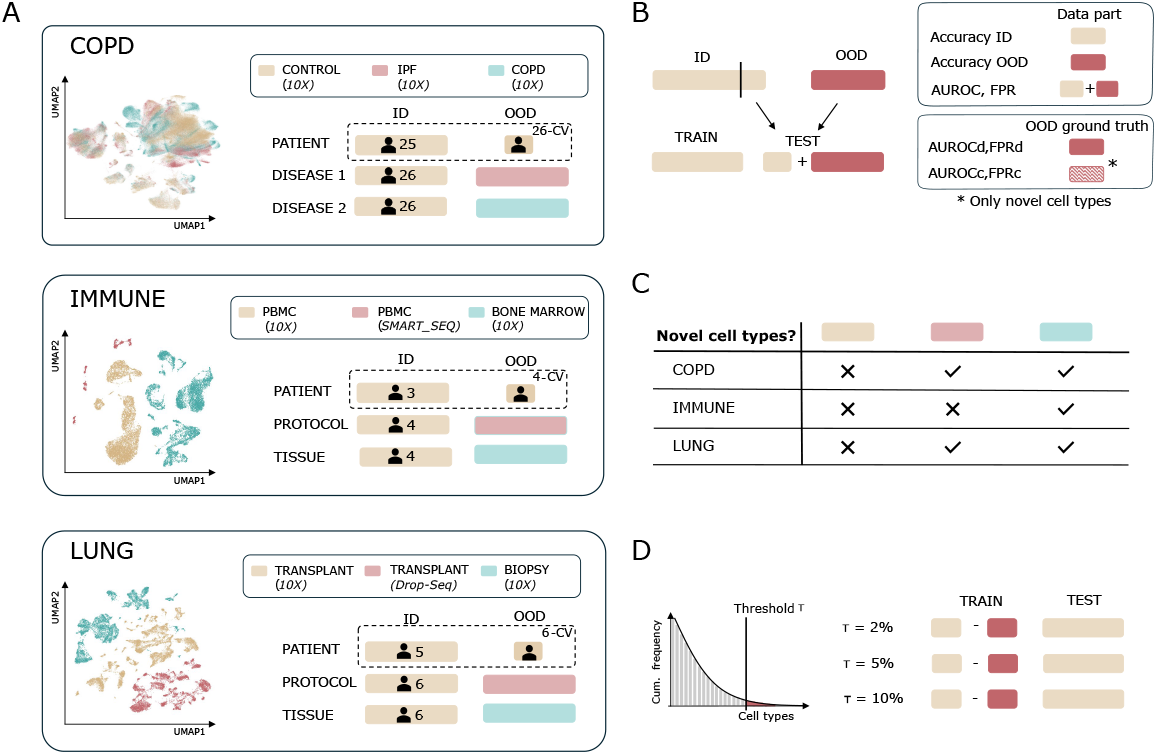
A graphical representation of **A**. the biological application settings where the six OOD methods are tested on, simulated with the Lung, Immune and COPD dataset Luecken et al. [2022], Adams et al. [2020] and **B**. the train-test split scheme for all the application settings that indicates which data parts are used to measure in-distribution and out-of-distribution performance (ID and OOD accuracy) and OOD detection evaluation (AUROC, FPR). For the OOD detection evaluation, which parts of the data are truly considered OOD (= the ground truth) for calculation of the metrics are also visualized. **C**. An indication of the presence of naturally occurring novel cell types in the OOD part of the test dataset for all three datasets. The colors correspond to the colors used in panel A to indicate the dataset parts. **D**. A visual representation of the minor novel cell type analyses, where for the 10X healthy control data of the COPD dataset Adams et al. [2020], a percentage of cell types, ranked based on their presence in the dataset, was excluded from the training data.

### 2.4 OOD scenarios

With the help of these datasets, we mimicked several application settings, where the test data could naturally contains a data shift. For the Immune and Lung dataset we considered the annotation of a new patient dataset, a dataset generated by a different protocol and a dataset originating from a different tissue. For the COPD dataset, again annotation of a new patient was considered, but also annotation of two dataset parts with a disease effect. Figure 1A visualizes for the three datasets the application settings and their corresponding ID and OOD data parts. To evaluate the effect of a patient shift across all the three datasets, a grouped cross-validation scheme was implemented so that each patient in the dataset was once considered as test data, the resulting metrics were averaged over all the patients (see Figure 1A). Patients with less than 500 sequenced cells in total were excluded from the analysis.

We evaluated annotation performance on the ID data and OOD data separately and evaluated OOD detection. In order to do this, a part of the ID data was included in the test data to evaluate the ID classification performance of the model, as illustrated in Figure 1B. Before we could evaluate OOD detection performance, a ground truth OOD label needed to be determined for all the samples in the test data, indicating whether each sample is ID or OOD. We evaluated OOD detection for two scenarios: (1) a scenario where a severe data shift is present, so the goal is to detect the entire OOD data part as OOD and (2) a scenario where the OOD detection goal is to detect novel cell types (cell types not present in the training data). This latter scenario is not applicable for all application settings as not all the test datasets (naturally) contain novel cell types (Figure 1C).

Next to these biological OOD settings, we also tested the OOD performance of the methods in an artificial OOD setting. For the 10X healthy control data of the COPD dataset, we excluded the 2,5 or 10 % least occurring cell types during training (see Figure 1D). This resulted in respectively the exclusion of 22, 34 and 42 out of 60 cell types, due to the severe imbalanced nature of the data. We will refer to this experiment as minor novel cell type detection in the rest of this manuscript. For all the training and test datasets (incl. the OOD data parts), with exception of the biological patient OOD setting, cell type populations with less than 10 observations were filtered out. For the biological patient OOD setting this filtering was performed on the entire dataset across all patients.

### 2.5 Evaluation metrics

To evaluate the cell type annotation performance, the ID accuracy and OOD accuracy metric are calculated based on the cell type labels assigned to the cells. The former is calculated on the part of the in-distribution data amended to the test dataset, the latter is calculated on the OOD data (see Figure 1B). For the OOD detection evaluation, the AUROC metric and the false positive rate (FPR) for a true positive rate (TPR) of 95% are reported based on the predicted OOD labels. To calculate these metrics, ground truth OOD labels needed to be assigned. To do so, we assigned a positive value (1) to OOD samples and a negative value (−1) to ID samples. We considered two scenarios (i) all samples from the OOD data part are OOD (AUROCd, FPRd) and (ii) only the samples with cell type labels unseen during training i.e. novel cell types (if present in the test data) are OOD (AUROCc, FPRc).

## 3 Results and discussion

### 3.1 OOD detection does not influence ID annotation

The results of the 6 OOD methods on the Immune and Lung datasets are presented in Tables A1 and A2, and Figure 2. The ID performance is high and similar across all methods, except for the Posterior Network. This was expected, as the OOD methods, with the exception of the Posterior Network, retain the main training objective to correctly classify ID cell types, similar to a general cell type annotation tool, and slightly adapt the learning loss or use the output of this classification task to perform OOD detection.

**Figure 2:**
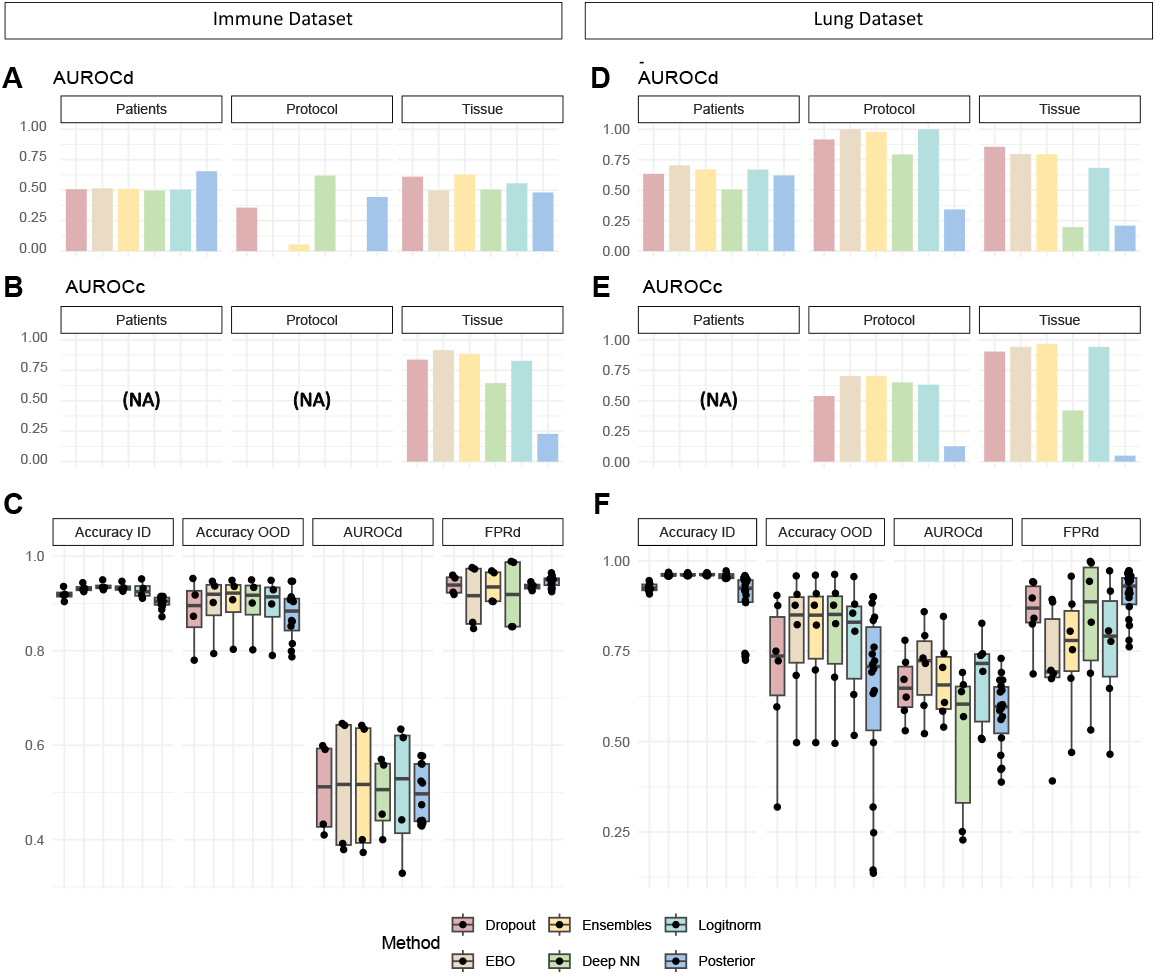
Overview of the OOD detection results of the 6 methods on the Immune (**A**., **B**. and **C**.) and the Lung dataset (**D**., **E**. and **F**.). Figures **A**. and **D**. display the AUROCd or AUROC dataset metric, where the AUROC is calculated when the entire OOD data part of the test dataset is desired to be detected as OOD, for all the 6 methods. Figures **B**. and **E**. display (if applicable) the AUROCc or AUROC cell type metric, where the AUROC metric is calculated when the goal is to detect naturally occurring novel cell types in the OOD data part of the test dataset, for all the 6 methods. The box plots in figures **C**. and **F**. show the ID accuracy, OOD accuracy, AUROCd and FPRd when the TPR is 95% for all the patients separately for the patient OOD splits. Every point represents one analysis, which for all methods besides the Posterior Network, corresponds with one patient.

The Posterior Network, however, performs significantly worse for ID classification in comparison to the other methods and gave unstable results across multiple analysis runs. To mitigate this instability, we ran all the Posterior Network analyses three times and reported the average. A possible explanation to this instability could be due to the loss function, as it has been reported that loss functions of deep evidential methods such as the Posterior Networks are unstable Bengs et al. [2022], Juergens et al. [2024]. The authors reported comparable results with other OOD methods for computer vision data, but single-cell data differs significantly, making it difficult to pinpoint the cause of the inferior performance. Notably, the authors themselves state that improving (ID) accuracy is not the goal of their method Charpentier et al. [2020].

Our hyperparameter tuning showed that non-linear networks consistently performed slightly better than linear ones, contradicting the literature Abdelaal et al. [2019], Köhler et al. [2021], Huang and Zhang [2021]. A recent paper suggested that non-linear methods outperform linear ones in a cross-tissue context with large-scale datasets Fischer et al. [2024]. However, our OOD patient splits occur within one tissue and with training datasets of a relative small size. Given the small performance differences and the reasonably non-complex datasets, we conclude that optimal ID classification performance may have been achieved for these datasets.

### 3.2 OOD methods are able to accurately detect novel cell types and possibly large data shifts

As mentioned before, we considered two possible OOD scenarios (i) the presence of a severe data shift resulting in (desired) OOD detection of the entire OOD data part and (ii) detection of novel cell types in the absence of a large data shift. In order to conclude which OOD scenario is desired for the different biological annotation applications, we calculated the Wasserstein’s distance across the first 100 principal components of the ID and OOD data parts. For more information on this calculation we refer the reader to the Appendix 1.3. Figure A2.A and B show these distances for each annotation application of the Lung and Immune datasets. The largest difference between the training and OOD distributions occurs in the protocol OOD scenario, indicating that OOD detection of the entire OOD protocol data part might be desired for both datasets.

Figure 2 shows the OOD detection performance of different methods for the Lung and Immune datasets. For the Immune dataset (Figure 2A-C), novel cell types naturally occurred only in the tissue OOD split, where the Energy-based OOD detection method performed best with an AUROC of 91.5%. In the protocol split, no method accurately detected the entire dataset as OOD, despite the large Wasserstein distances (Figure A2.B). For the Lung dataset (Figure 2D- F), most methods accurately detected the entire protocol OOD data part, with Energy-based OOD detection achieving an AUROC of 100%. The naturally occurring novel cell types in the tissue split were best recognized by Ensembles (AUROC 96.%), followed closely by the Energy-based OOD (AUROC 94.2%) and Logitnorm (AUROC 94.1%) methods. Overall, the methods performed well in recognizing novel cell types in the tissue OOD split, with the Energy-based OOD method being the top performer.

Based on the mixed results across the two datasets for the protocol splits, it is unclear if the methods can accurately flag entire data parts affected by large batch effects due to protocol differences or other variations leading to covariate shifts. The Immune dataset, which has the largest distance between protocol distributions, is not flagged, while the Lung OOD protocol is. Given the clear distinctions in the UMAPs in Figure 1, we believe separation should be possible, and these protocols should be able to detect the splits based on their mode of operation. A potential bottleneck is that the OOD methods operate in the embedded space of the ID annotation task, which may be suboptimal for OOD detection, especially for detecting covariate shifts.

We also visualized for the patient splits the relation between the OOD Accuracy and AUROCd in Figure A3, to see if severe patients shifts are being picked up by the best performing methods: Ensembles and EBO. It seems that most of patients with a severe data shift are picked up as being OOD by the detection methods across the datasets. So overall, the OOD methods are able to pick up severe data shifts, though not consistently across all datasets.

### 3.3 Patient, protocol and tissue each have a clear effect on annotation with increasing severity

A key question is how the biological settings affect the ID and OOD annotation performance? Based on the results reported in the Tables A1 and A2, a conclusion could be made that, on average, patient effects are negligible as the accuracy drop from ID to OOD annotation is relatively small, especially for the Immune dataset. However, as visualized in Figure 2C and F, for some patients a clear data shift is occurring, indicating the importance of checking or accounting for possible patient effects during cell type annotation.

To find out the influence of tissue and protocol shifts on cell type annotation, we re-visualized the results in Figures A4 and A5 to clearly see the influence of the different OOD splits. These results show that both a protocol and tissue shift have a severe influence on annotation performance and that the influence of a new tissue is significantly larger than that of a new protocol. These results contradict the results of the Wasserstein distance measures. An explanation for this lies in the calculation of the Wasserstein distance and the nature of the data generated by different protocols. Protocols like 10X, SMART-Seq, and Drop-seq have varying sensitivities, i.e. detected genes per cell, and capture efficiency, leading to noticeable global data shifts captured by the Wasserstein distance metric in the PCA space Mereu et al. [2020], Ziegenhain et al. [2017]. However, cell type annotation mainly relies on a limited set of marker genes. If the signal in the individual marker genes is conserved, cell type annotation won’t be significantly affected by the large data shifts. Data distributions of different tissues sequenced with the same protocol will show biological variation and data shifts, which may be present in marker genes. This can hinder cell type annotation without greatly impacting the Wasserstein metric. Therefore, a large Wasserstein distance will indicate a severe data shift, that ideally would be flagged by the OOD methods. But the Wasserstein distance does not necessarily reflect annotation performance on the OOD dataset.

### 3.4 Integration increases annotation performance but might hinder OOD detection

The results for the integrated COPD dataset are presented in Table A3 and Figures A1 and A6. Integration significantly improves the annotation performance of the OOD data. The performance drops between ID and OOD annotation performance seen on the Lung and Immune dataset are not visible on this dataset. For this dataset, the COPD disease effect seems the most compensated, the ID and OOD accuracy difference is the smallest for this set-up. The drop increases slightly for the IPF data and between the patients. However, the detection performance of naturally occurring novel cell types also drops. For the IPF OOD data part, Ensembles perform the best with an AUROC of 75.6%. For the COPD OOD data part Energy-based OOD performs the best with an AUROC of 70.5%. Although these results are not bad, they are not as strong as before. It is difficult to determine if this performance drop is solely due to integration, as the COPD dataset is more complex with more novel cell types. Based on these results, we can conclude that Energy-based OOD was the most effective method for novel cell type detection in our evaluation across the three datasets. Additionally, it calculates a post-inference OOD score, making it computationally efficient.

For this dataset, we did not report the results of Posterior Networks for the COPD datasets, since the method failed to learn anything during training across all our setups (OOD scenarios, network configurations, etc.) i.e. the uncertainty-aware loss value did not change during training. To address this, we tried varying the learning rates (1e-1 to 1e-8), we increased the number of epochs and relaxed the early stopping criteria, but no improvement was observed. Since Posterior Networks train a Normalizing Flow and embedding network in one pass, identifying the bottleneck is challenging. Given the COPD dataset’s larger size, more features, and more classes compared to the other datasets (see Table 2), there could be multiple reasons why this method worked on other datasets but not on the COPD dataset. We hypothesize that the reason for the network’s inability to learn from this dataset lies in challenges associated with density estimation; however, further investigation regarding the obtained results are required and is left as future work.

**Table 2:** The size, number of label classes (cell type populations) and the different data splits (experimental setups) used to evaluate OOD for the three considered datasets (the Immune, Lung and COPD) dataset.

| Dataset | N° cells<br>ID/OOD | N° features | N° cell populations<br>ID/OOD (NC)<sup>1</sup> | Experimental setups |  |  |
| --- | --- | --- | --- | --- | --- | --- |
|  |  |  |  | Split | Protocol | Tissue / Disease |
| **Immune** | 8,829<sup>2</sup> | 12,303 | 10<sup>2</sup> | Patient | 10X | PBMC |
|  | 8,829/1,022 | 12,303 | 10/10 (-) | Protocol | Smart-seq | PBMC |
|  | 8,829/9,581 | 12,303 | 10/16 (6) | Tissue | 10X | Bone Marrow |
| **Lung** | 11,828<sup>2</sup> | 15,148 | 13<sup>2</sup> | Patient | 10X | Transplant |
|  | 12,716/9,701 | 15,148 | 13/11 (1) | Protocol | Drop-seq | Transplant |
|  | 12,716/10,033 | 14,148 | 13/10 (3) | Tissue | 10X | Biopsy |
| **COPD** | 95,506<sup>2</sup> | 45 497 | 78<sup>2</sup> | Patient | 10X | Control |
|  | 96,260/147,158 | 45,947 | 78 / 97 (19) | Disease 1 | 10X | IPF |
|  | 92,260/69,423 | 45,947 | 78 / 86 (8) | Disease 2 | 10X | COPD |

<sup>1</sup>NC stands for new cell types, cell types not present in the ID data.
<sup>2</sup>As for the patient splits a group cross-validation scheme is used, where each patient is separately used as test data, the total number of cells and cell type populations across all patients is reported.

We also performed a synthetic minor novel cell type split on the COPD 10X control data (Figure A1B and A7). Here, not Energy-based OOD, but Deep NN was the best performer, underlining the importance of evaluating OOD methods in real-life scenarios. For the 2%, 5%, and 10% hold-out schemes the AUROC values for the novel cell type detection were 82.8 86.5%, and 80.1%, respectively. This demonstrates the method’s insensitivity to the number of novel cell types, as is the case for all the other methods, with exception of the Ensembles, where detection performance slightly improved as more cell types were rejected.

## 4 Conclusion

In this paper, we evaluated six established OOD detection methods from the computer vision field for cell type annotation of single-cell transcriptomics data, specifically for the detection of data shifts and novel cell types. We evaluated their performance on a synthetic use-case and on real-life biological data shifts such as the introduction of new patients, a new protocol, a new tissue or a new disease. We performed these analyses on two normal single-cell datasets and one integrated dataset. Based on our results, we recommend to use Energy-based OOD detection for novel cell type detection, as it overall performed best in our evaluation on real-life biological data shifts. Energy-based OOD is also computationally very efficient and easy to implement as it uses a post-inference score for OOD detection.We illustrated the importance of including real-life biological scenario’s in the OOD detection evaluation as they had severe impact on the cell type annotation performance and gave different results for OOD detection in comparison to the synthetic evaluation. Our results showed promise for the OOD detection methods to also be used for detecting data shifts, but the results were inconsistent across dataset, so more research needs to be performed before a firm conclusion can be made. Lastly, we saw that integration does increase cell type annotation on data under the influence of a data shift, as expected. But integration might also negatively influence novel cell type detection.

## 6 Data availability

The datasets were derived from sources in the public domain, they can be accessed through the repositories mentioned in the corresponding articles. Code to reproduce the analyses in this paper is freely available at: https://github.com/Latheuni/OODeval.git

## 8 Author contributions statement

L.T, T.M, Y.S and W.W. conceived the experiments; L.T. conducted the experiments, analyzed the results and wrote the manuscript; L.T, T.M, Y.S and W.W. reviewed the manuscript.

## Footnotes

1 A proper scoring rule *S*_*proper*_ is a scoring rule such that *S*_*proper*_(*P*_*θ*_, *P*) ≤ *S*_*proper*_(*P, P*) if and only if *P*_*θ*_(*y*|**x**) = *P* (*y*|**x**).

