## Appendix for "Evaluation of out-of-distribution detection methods for data shifts in single-cell transcriptomics"

### Appendix 1: Extra information

#### A1.1 Dataset information and preprocessing

We evaluated the out-of-distribution methods across 3 different datasets: the Lung, Immune and COPD dataset [Luecken et al., 2022, Adams et al., 2020]. We obtained the (preprocessed) Lung and Immune datasets from Luecken et al. (2022) Luecken et al. [2022], a single-cell RNA-seq integration benchmark where multiple open source datasets were assembled into new datasets with different protocols, tissues or disease states and a homogenized annotation[1]. The Lung dataset consists of three healthy 10X transplant datasets, one Drop-seq transplant dataset and one 10X lung biopsy dataset. The samples from the biopsy dataset were extracted from the airways, while the transplant samples were taken from the parenchyma. Luecken et al. (2022) Luecken et al. [2022] performed quality control, scran pooling Lun et al. [2016] on the individual datasets and harmonized the annotations. The (integrated) Immune dataset contains peripheral blood mononuclear cell (PBMC) data, sequenced with SMART-seq and 10X, together with Bone marrow data sequenced with 10X. For this data, Luecken et al. (2022) Luecken et al. [2022] performed quality control on all data parts and scran pooling Lun et al. [2016] for the 10X data. Both datasets were log(1+x) transformed. The COPD dataset was generated by Adams et al. (2020) Adams et al. [2020] and contains samples from healthy (smoker and non-smoker) patients, patients with idiopathic pulmonary fibrosis (IPF) and patients with chronic obstructive pulmonary disease (COPD), all sequenced with 10X. We normalized the count data with a scale factor of 10,000 and performed log(1+x) transformation, similar to Adams et al. (2020) Adams et al. [2020]. Characteristics of the datasets can be found in Table 2 in the main manuscript.

#### A1.2 Model construction

Each OOD method was implemented with multiple NN architectures for $f$. For $f$, fully-connected neural networks with 0,2,5 and 10 hidden layers with a ReLU activation were implemented, resulting in one linear and three non-linear network architectures. For the non-linear NN, i.e. those with hidden layers, the hidden layers of the NN architecture each consisted out of 50, 100 or 1000 nodes. Early stopping was implemented during training and a max training epoch limit was set to 350. The used batch size during training was 254, the learning rate was set to 1e-4. The results of the best overall performing NN, based on the best scoring ID Accuracy and Balanced Accuracy performance were reported.

The training dataset was split into a validation dataset for early stopping with a size of 20%. In order to evaluate ID annotation performance, a part of the training dataset, of equal size to the validation dataset, was splitted of and concatenated to the test dataset (See Figure 1B) to evaluate in-distribution performance was set at the same size as the validation dataset. For the minor cell type detection, this part makes up the entire test dataset.

For all the methods, beside the Posterior Networks and Deep NN method, the default values of the hyperparameters were used i.e. those recommended by the original authors. For the Posterior Network, we used an Inverse autoregressive Flow Kingma et al. [2016] and tested the method with 8,10 and 20 number of transformations and a regularization factor in the Bayesian loss of 1e-2 and 1e-4. The use of Posterior Networks, trained with the uncertainty-aware loss, resulted in very unstable performance with regard to our chosen evaluation metrics. As a result, to make a fair comparison to the other methods, these models configurations were each independently trained and tested 3 times and the resulting metrics were averaged. For the Deep NN method, the tested numbers of nearest neighbors considered during the score calculation were 50, 100, 200, 500 and 1000.

#### A1.3 Wasserstein metric

In order to conclude which OOD scenario is desired for the different biological annotation applications, we constructed a PCA with 100 components on the ID dataset. We then transformed the OOD data to this constructed PCA space and calculated the Wasserstein distance between each principal component of the transformed ID and OOD data. We visualized these distances in Figure A2, where each point corresponds to one distance value.

#### Appendix 2: Tables and Figures

**Table A1.** Results of the Lung dataset, for the six methods (LogitNorm, Dropout, Ensembles, Energy-based OOD ( EBO), Deep NN and Posterior Networks) across the three OOD data splits (Patients, Protocol and Tissue). The ID and OOD cell type annotation accuracy is reported (see Figure 1B) together with the AUROCd/c and FPRd/c, which is the FPR when the TPR is 95%, for OOD detection. The d/c stands for dataset or celltype and refers to which part of the test dataset is set as ground-truth OOD: resp. the OOD data part of the test dataset or the naturally occuring cell types in the OOD data part of the test dataset.

| | | Lung dataset | | | | | |
|---|---|---|---|---|---|---|---|
| Setup | Metric | LogitNorm | Dropout | Ensembles | EBO | Deep NN | Posterior[1] |
| Patients[2] | Accuracy ID | 0.958(0.008) | 0.948(0.008) | 0.962(0.003) | 0.961(0.004) | 0.962(0.003) | 0.926(0.01) |
| | Accuracy OOD | 0.774(00152) | 0.757(0.139) | 0.794(0.156) | 0.791(0.157) | 0.791(0.159) | 0.705(0.162) |
| | Balan. Acc. ID | 0.883(0.046) | 0.752(0.139) | 0.874(0.05) | 0.872(0.048) | 0.874(0.055) | 0.692(0.032) |
| | Balan. Acc. OOD | 0.85(0.056) | 0.791(0.076) | 0.853(0.059) | 0.848(0.059) | 0.853(0.06) | 0.735(0.054) |
| | Accuracy Drop | 0.184(0.152) | 0.191(0.139) | 0.168(0.156) | 0.17(0.157) | 0.171(0 159) | 0.221(0.152) |
| | $AUROC_{d/c}$ | 0.669(0.121)/- | 0.634(0.103)/- | 0.671(0.105)/- | **0.704(0.113)**/- | 0.506(0.192)/- | 0.544(0.109)/- |
| | $FPR_{d/c}$ | 0.764(0.169)/- | 0.856(0.125)/- | 0.757(0.156)/- | **0.705(0.167)**/- | 0.831(0.172)/- | 0.929(0.029)/- |
| Protocol | Accuracy ID | 0.954 | 0.943 | 0.959 | 0.957 | 0.956 | 0.93 |
| | Accuracy OOD | 0.618 | 0.571 | 0.628 | 0.632 | 0.64 | 0.46 |
| | Balan. Acc. ID | 0.847 | 0.723 | 0.844 | 0.881 | 0.839 | 0.74 |
| | Balan. Acc. OOD | 0.656 | 0.567 | 0.665 | 0.667 | 0.679 | 0.368 |
| | Accuracy Drop | 0.336 | 0.372 | 0.331 | 0.325 | 0.316 | 0.47 |
| | $AUROC_{d/c}$ | 0.999/0.632 | 0.916/0.539 | 0.978/0.704 | **1.0/0.706** | 0.793/0.651 | 0.512/0.407 |
| | $FPR_{d/c}$ | 0.006/0.649 | 0.316/0.777 | 0.079/**0.582** | **0.001**/0.601 | 0.583/0.661 | 0.947/0.844 |
| Tissue | Accuracy ID | 0.955 | 0.945 | 0.957 | 0.956 | 0.955 | 0.538 |
| | Accuracy OOD | 0.335 | 0.305 | 0.312 | 0.314 | 0.313 | 0.102 |
| | Balan. Acc. ID | 0.842 | 0.768 | 0.843 | 0.845 | 0.846 | 0.306 |
| | Balan. Acc. OOD | 0.571 | 0.464 | 0.561 | 0.547 | 0.56 | 0.252 |
| | Accuracy Drop | 0.62 | 0.64 | 0.645 | 0.642 | 0.642 | 0.436 |
| | $AUROC_{d/c}$ | 0.683/0.941 | **0.856**/0.903 | 0.795/**0.967** | 0.769/0.942 | 0.197/0.42 | 0.202/0.202 |
| | $FPR_{d/c}$ | 0.999/0.164 | **0.607**/0.283 | 0.71/**0.122** | 0.749/0.17 | 1.0/0.787 | 0.976/0.932 |

[1]Due to the unstability of the method, we ran this analysis three times, the average metric value of the three runs is reported.

[2]For the patient scenario the average metric value over the patients is reported together with the standard deviation in between brackets

**Table A2.** Results of the Immune dataset, for the six methods (LogitNorm, Dropout, Ensembles, Energy-based OOD ( EBO), Deep NN and Posterior Networks) across the three OOD data splits (Patients, Protocol and Tissue). The ID and OOD cell type annotation accuracy is reported (see Figure 1B) together with the AUROCd/c and FPRd/c, which is the FPR when the TPR is 95%, for OOD detection. The d/c stands for dataset or celltype and refers to which part of the test dataset is set as ground-truth OOD: resp. the OOD data part of the test dataset or the naturally occuring cell types in the OOD data part of the test dataset.

| | | Immune dataset | | | | | |
|---|---|---|---|---|---|---|---|
| Setup | Metric | LogitNorm | Dropout | Ensembles | EBO | Deep NN | Posterior[1] |
| Patients[2] | Accuracy ID | 0.928 (0.016) | 0.919 (0.011) | 0.928 (0.016) | 0.932 (0.007) | 0.934(0.008) | 0.905(0.009) |
| | Accuracy OOD | 0.892 (0.061) | 0.881 (0.065) | 0.892 (0.061) | 0.895 (0.061) | 0.897(0.057) | 0.87(0.046) |
| | Balan. Acc. ID | 0.848 (0.054) | 0.611 (0.03) | 0.848 (0.054) | 0.828 (0.043) | 0.855 (0.054) | 0.617(0.029) |
| | Balan. Acc. OOD | 0.817 (0.063) | 0.652 (0.034) | 0.817 (0.063) | 0.79 (0.058) | 0.81 (0.042) | 0.673(0.041) |
| | Accuracy Drop | 0.036 (0.063) | 0.038 (0.066) | 0.036 (0.063) | 0.037 (0.061) | 0.037 (0.058) | 0.035(0.047) |
| | AUROCd/AUROCc | 0.505 (0.126)/- | 0.508 (0.087)/- | 0.505 (0.126)/- | **0.515 (0.129)**/- | 0.496 (0.071)/- | 0.502(0.055) |
| | FPRd/FPRc | 0.936 (0.007)/- | 0.939 (0.018)/- | 0.936 (0.007)/- | **0.914 (0.061)**/- | 0.92 (0.069)/- | 0.94(0.011) |
| Protocol | Accuracy ID | 0.925 | 0.92 | 0.935 | 0.934 | 0.934 | 0.597 |
| | Accuracy OOD | 0.75 | 0.263 | 0.785 | 0.733 | 0.774 | 0.283 |
| | Balan. Acc. ID | 0.862 | 0.629 | 0.867 | 0.783 | 0.862 | 0.228 |
| | Balan. Acc. OOD | 0.807 | 0.43 | 0.868 | 0.832 | 0.86 | 0.436 |
| | Accuracy Drop | 0.175 | 0.657 | 0.15 | 0.201 | 0.16 | 0.314 |
| | AUROCd/AUROCc | 0.0/- | 0.357/- | 0.055/- | 0.0/- | .62/- | **0.935**/- |
| | FPRd/FPRc | 1.0/- | 1.0/- | 1.0/- | 1.0/- | 0.518/- | **0.393/-** |
| Tissue | Accuracy ID | 0.936 | 0.913 | 0.936 | 0.935 | 0.933 | 0.901 |
| | Accuracy OOD | 0.517 | 0.5 | 0.539 | 0.538 | 0.53 | 0.502 |
| | Balan. Acc. ID | 0.837 | 0.609 | 0.878 | 0.876 | 0.847 | 0.638 |
| | Balan. Acc. OOD | 0.441 | 0.256 | 0.483 | 0.482 | 0.468 | 0.388 |
| | Accuracy Drop | 0.419 | 0.413 | 0.397 | 0.397 | 0.403 | 0.399 |
| | AUROCd/AUROCc | 0.556/0.827 | 0.611/0.837 | **0.629**/0.884 | 0.497/**0.915** | 0.547/0.644 | 0.632/0.757 |
| | FPRd/FPRc | 1.0/0.623 | 0.978/0.562 | 0.99/0.387 | 1.0/**0.377** | **0.896**/0.58 | 0.963/0.727 |

[1]Due to the unstability of the method, we ran this analysis three times, the average metric value of the three runs is reported.

[2]For the patient scenario the average metric value over the patients is reported together with the standard deviation in between brackets

**Table A3.** Results of the COPD dataset, for the six methods (LogitNorm, Dropout, Ensembles, Energy-based OOD ( EBO), Deep NN and Posterior Networks) across the three biological OOD data splits (Patients, Disease1 and Disease2) and the minor novel cell type annotation task, where the 2,5 and 10% least occuring cell type populations were excluded during training. The ID and OOD cell type annotation accuracy is reported (see Figure 1B) together with the AUROCd/c and FPRd/c, which is the FPR when the TPR is 95%, for OOD detection. The d/c stands for dataset or celltype and refers to which part of the test dataset is set as ground-truth OOD: resp. the OOD data part of the test dataset or the naturally occuring cell types in the OOD data part of the test dataset.

| **OOD scenario** | Metric | COPD dataset | | | | | |
|---|---|---|---|---|---|---|---|
| | | LogitNorm | Dropout | Ensembles | EBO | Deep NN | Posterior[1] |
| **Patients[2]** | Accuracy ID | 0.871 (0.007) | 0.849 (0.01) | 0.889 (0.002) | 0.884 (0.003) | 0.883 (0.003) | - |
| | Accuracy OOD | 0.771 (0.161) | 0.725 (0.213) | 0.789 (0.155) | 0.786 (0.154) | 0.781 (0.166) | - |
| | BAcc ID | 0.589 (0.04) | 0.349 (0.046) | 0.634 (0.015) | 0.623 (0.022) | 0.619 (0.025) | - |
| | BAcc OOD | 0.728 (0.09) | 0.617 (0.128) | 0.749 (0.079) | 0.749 (0.079) | 0.743 (0.082) | - |
| | Accuracy Drop | 0.1 (0.161) | 0.124 (0.213) | 0.1 (0.155) | 0.089 (0.154) | 0.102 (0.134) | - |
| | $AUROC_{d/c}$ | 0.614 (0.105)/- | 0.599 (0.094)/- | 0.598 (0.115)/- | **0.622 (0.102)**/- | 0.427 (0.103)/- | - |
| | $FPR_{d/c}$ | 0.82 (0.097)/- | 0.857 (0.079)/- | 0.846 (0.108)/- | **0.807 (0.121)**/- | 0.924 (0.085)/- | - |
| **Disease 1** | Accuracy ID | 0.875 | 0.843 | 0.884 | 0.884 | 0.877 | - |
| | Accuracy OOD | 0.824 | 0.732 | 0.835 | 0.838 | 0.819 | - |
| | BAcc ID | 0.66 | 0.322 | 0.624 | 0.673 | 0.606 | - |
| | BAcc OOD | 0.476 | 0.217 | 0.445 | 0.483 | 0.433 | - |
| | Accuracy Drop | 0.051 | 0.111 | 0.049 | 0.046 | 0.058 | |
| | $AUROC_{d/c}$ | **0.635**/0.689 | 0.606/0.726 | 0.567/**0.756** | 0.618/0.632 | 0.365/0.71 | - |
| | $FPR_{d/c}$ | **0.888**/0.962 | 0.904/0.883 | 0.916/**0.838** | 0.906/0.96 | 0.987/0.965 | - |
| **Disease 2** | Accuracy ID | 0.875 | 0.839 | 0.884 | 0.885 | 0.879 | - |
| | Accuracy OOD | 0.829 | 0.766 | 0.846 | 0.843 | 0.839 | - |
| | Accuracy drop | 0.046 | 0.076 | 0.038 | 0.042 | 0.04 | - |
| | BAcc ID | 0.589 | 0.314 | 0.616 | 0.646 | 0.606 | - |
| | BAcc OOD | 0.47 | 0.24 | 0.496 | 0.516 | 0.433 | - |
| | Accuracy Drop | 0.046 | 0.076 | 0.038 | 0.042 | 0.04 | |
| | $AUROC_{d/c}$ | **0.612**/0.606 | 0.563/0.603 | 0.55/0.615 | **0.612/0.705** | 0.394/0.677 | - |
| | $FPR_{d/c}$ | 0.925/0.989 | 0.918/0.904 | **0.893/0.868** | 0.899/0.878 | 0.979/0.945 | - |
| **2%** | Accuracy ID | 0.894 | 0.856 | 0.901 | 0.895 | 0.9 | - |
| | Accuracy OOD | 0.0 | 0.0 | 0.0 | 0.0 | 0.0 | - |
| | BAcc ID | 0.851 | 0.673 | 0.853 | 0.848 | 0.871 | - |
| | BAcc OOD | 0.0 | 0.0 | 0.0 | 0.0 | 0.0 | - |
| | AUROC | 0.708 | 0.754 | 0.719 | 0.748 | **0.828** | - |
| | FPR | 0.901 | 0.854 | 0.818 | 0.862 | **0.645** | - |
| **5%** | Accuracy ID | 0.898 | 0.893 | 0.909 | 0.903 | 0.905 | - |
| | Accuracy OOD | 0.0 | 0.0 | 0.0 | 0.0 | 0.0 | - |
| | BAcc ID | 0.866 | 0.868 | 0.895 | 0.879 | 0.884 | - |
| | BAcc OOD | 0.0 | 0.0 | 0.0 | 0.0 | 0.0 | - |
| | AUROC | 0.801 | 0.681 | 0.675 | 0.748 | **0.865** | - |
| | FPR | 0.739 | 0.856 | 0.816 | 0.833 | **0.547** | - |
| **10%** | Accuracy ID | 0.883 | 0.904 | 0.914 | 0.909 | 0.906 | - |
| | Accuracy OOD | 0.0 | 0.0 | 0.0 | 0.0 | 0.0 | - |
| | BAcc ID | 0.86 | 0.901 | 0.92 | 0.921 | 0.915 | - |
| | BAcc OOD | 0.0 | 0.0 | 0.0 | 0.0 | 0.0 | - |
| | AUROC | 0.688 | 0.659 | 0.687 | 0.711 | **0.801** | - |
| | FPR | 0.862 | 0.892 | 0.829 | 0.809 | **0.605** | - |

[1]As mentioned in the main text, we had problems getting the Posterior Network to learn on the COPD dataset. After trying various solutions and still failing, we decided to not report the results for the Posterior network numerically, as these results can't be trusted to be reproducible.

[2]For the patient scenario the average metric value over the patients is reported together with the standard deviation in between brackets.

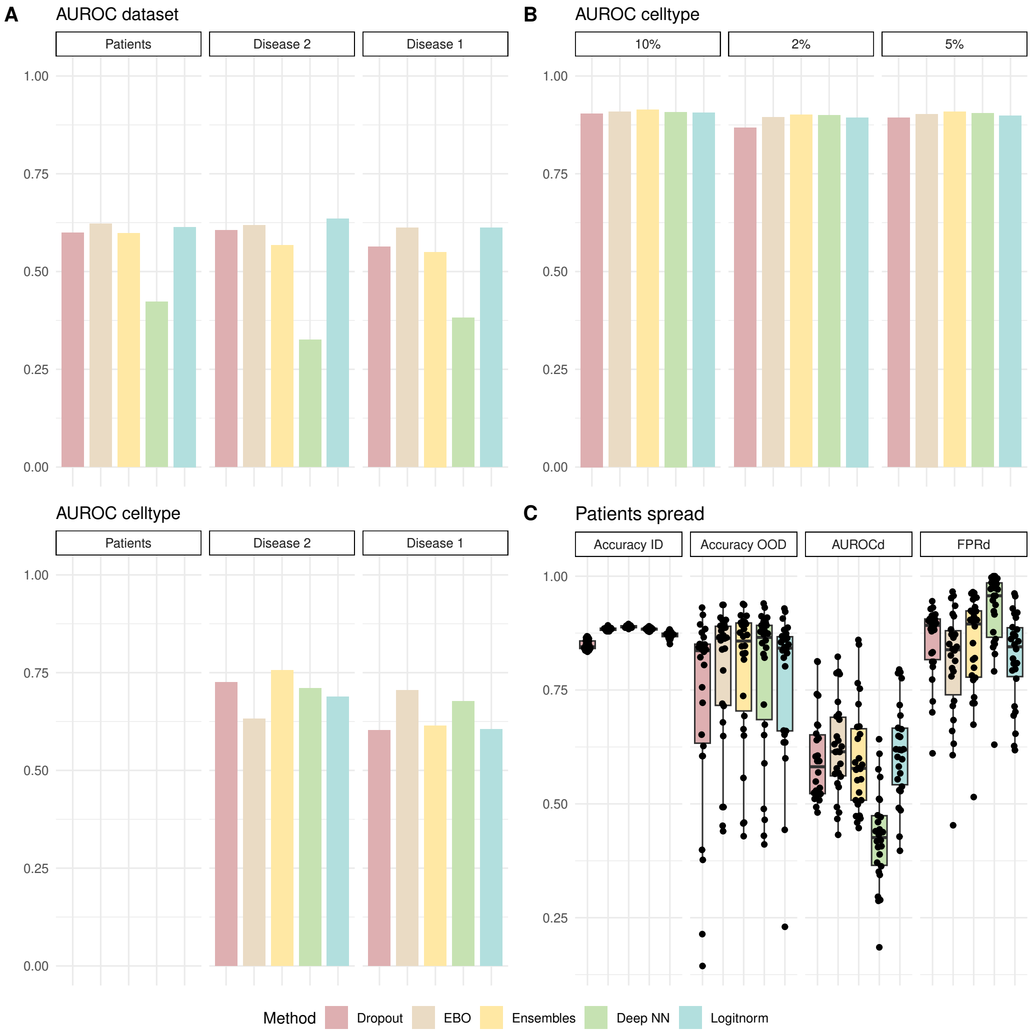


**Figure A1.** Overview of the OOD detection results for the COPD dataset. **A.** visualizes the AUROCd or AUROC dataset metric (top) and AUROCc or AUROC celltype metric for the 6 methods. The former metric corresponds to the AUROC metric calculated for the OOD detection problem when the entire OOD data part of the test dataset is desired to be detected as OOD. The latter metric is calculates when the goal is to detect naturally occuring novel cell types in the OOD data part of the test dataset. **B.** visualizes the AUROC metric for the minor novel celltype detection. Here, the 2,5 and 10% least occuring cell type groups are excluded during training, and need to be detected by the methods in the test dataset.**C.** shows for the patient OOD split, the ID accuracy, OOD accuracy, AUROCd and FPRd when the TPR is 95% for all

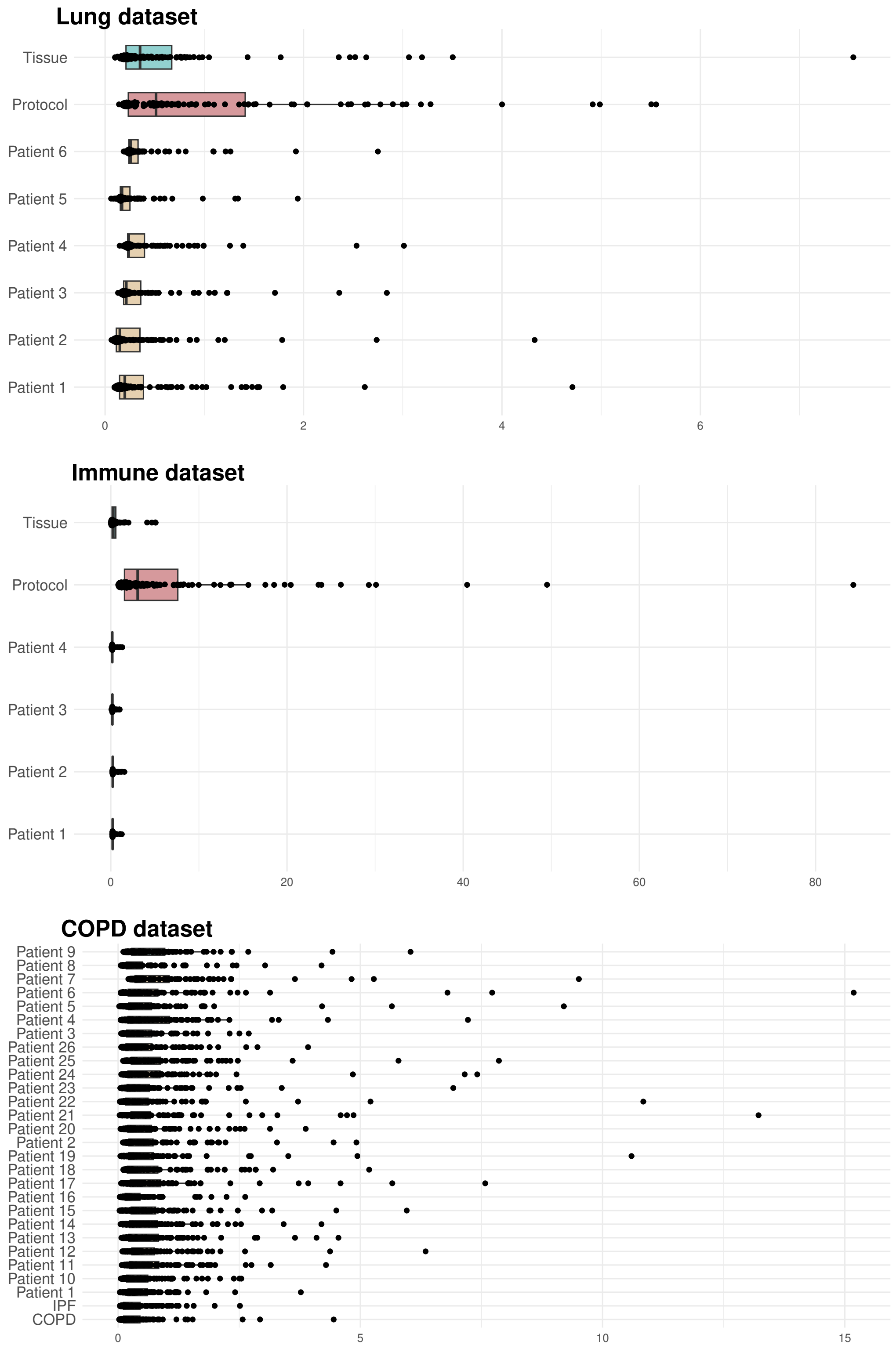


**Figure A2.** Visualization of the Wasserstein distances calculated between the first 100 PCA's of the training dataset and OOD data part of the test dataset across al the dataset splits for all three datasets. Each point represents one distance value.

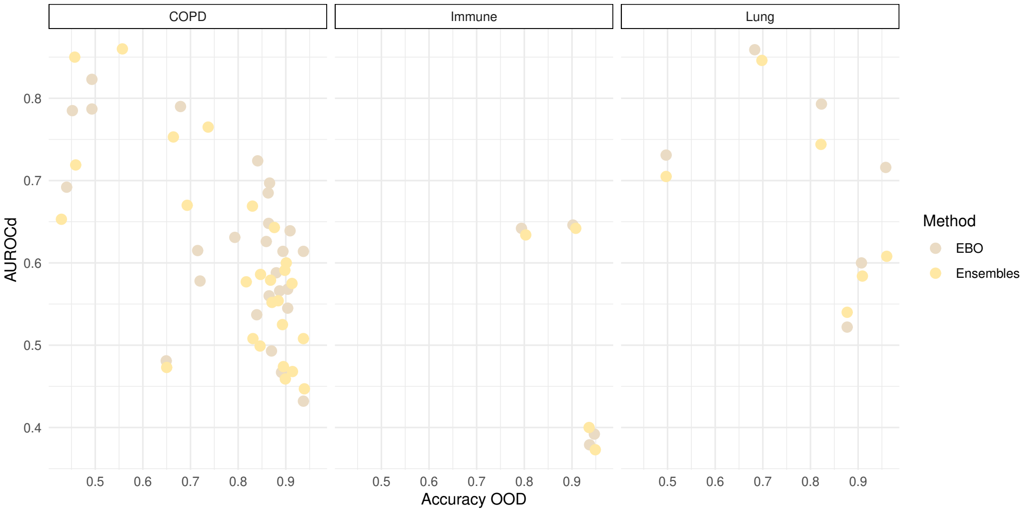


**Figure A3.** Scatter plot of the AUROCd and OOD accuracy of the Ensembles and Energy-based OOD method for the Lung, Immune and COPD dataset.

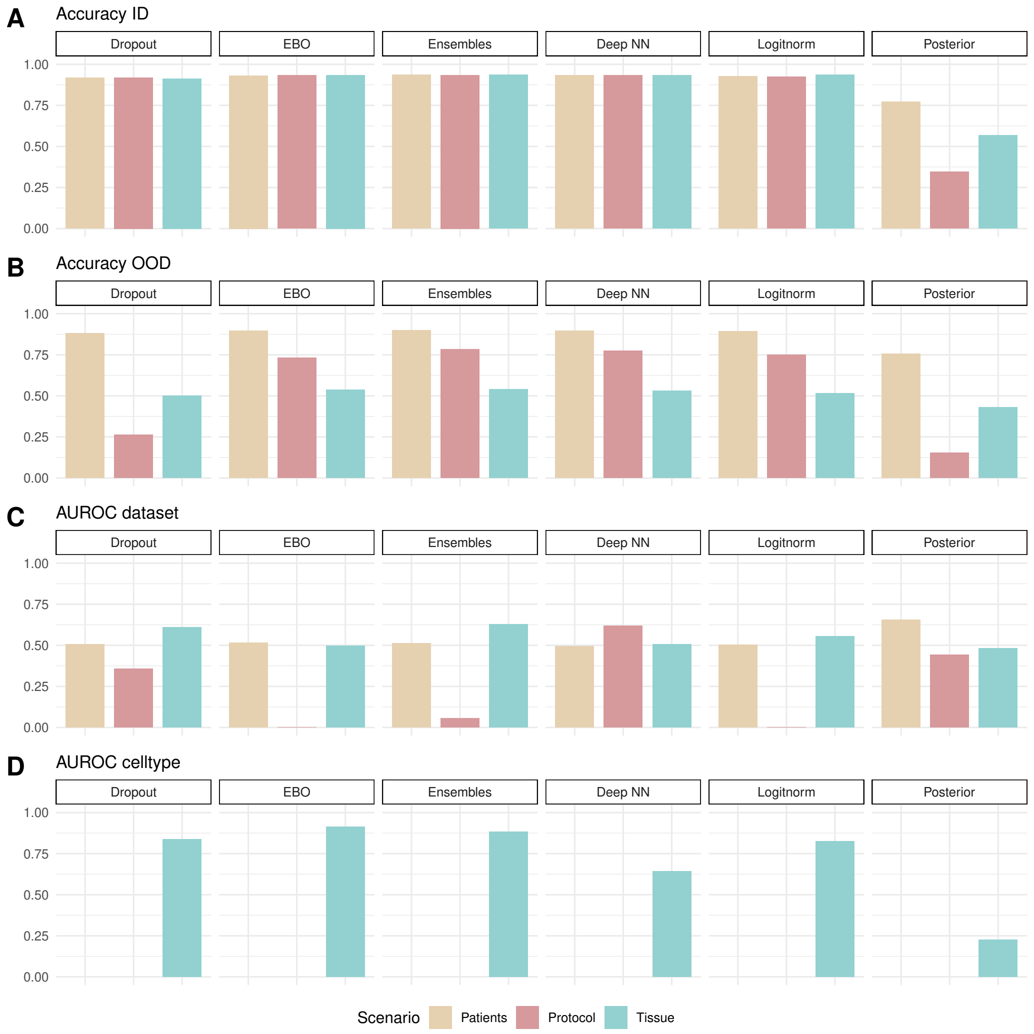


**Figure A4.** Visualisation of the ID annotation and OOD annotation results of the Immune dataset, for every method across the data splits (for patient splits the overall average is reported).

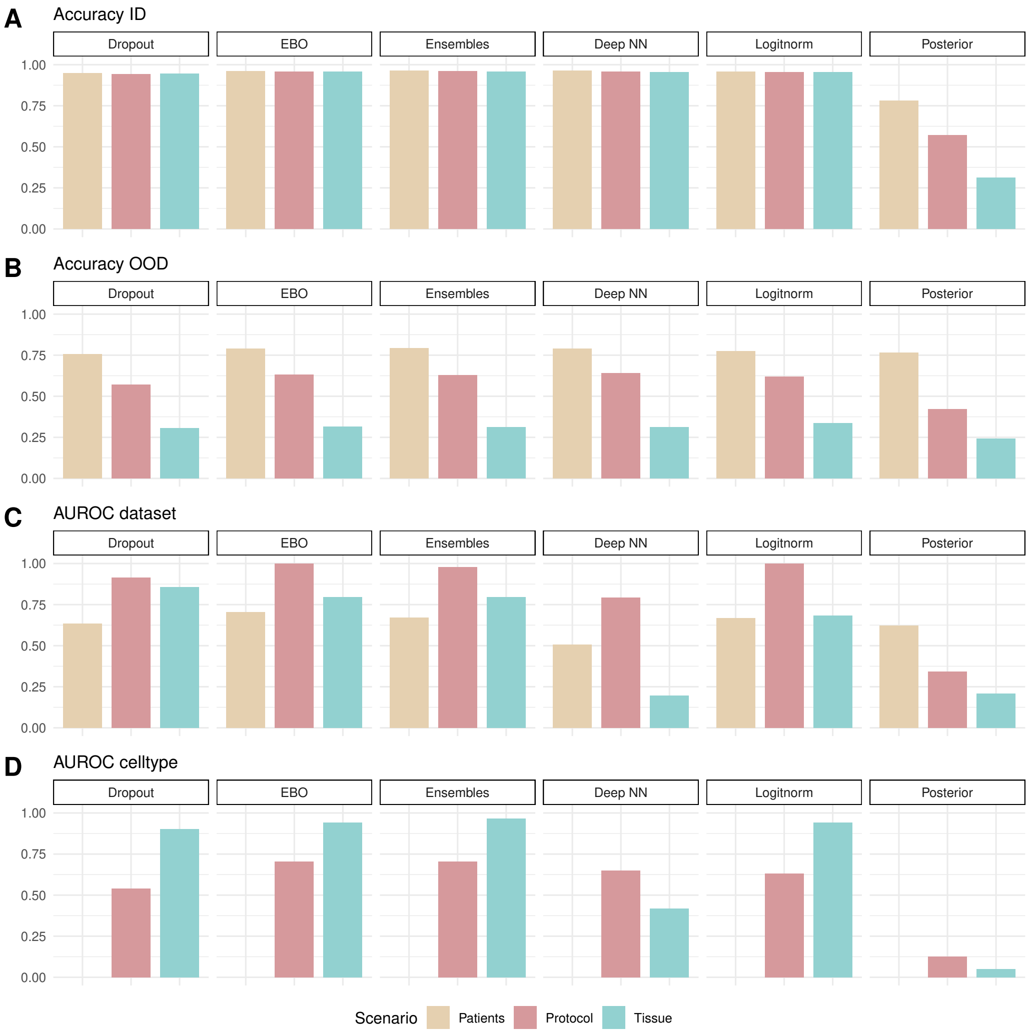


**Figure A5.** Visualisation of the ID annotation and OOD annotation results of the Lung dataset, for every method across the data splits (for patient splits the overall average is reported).

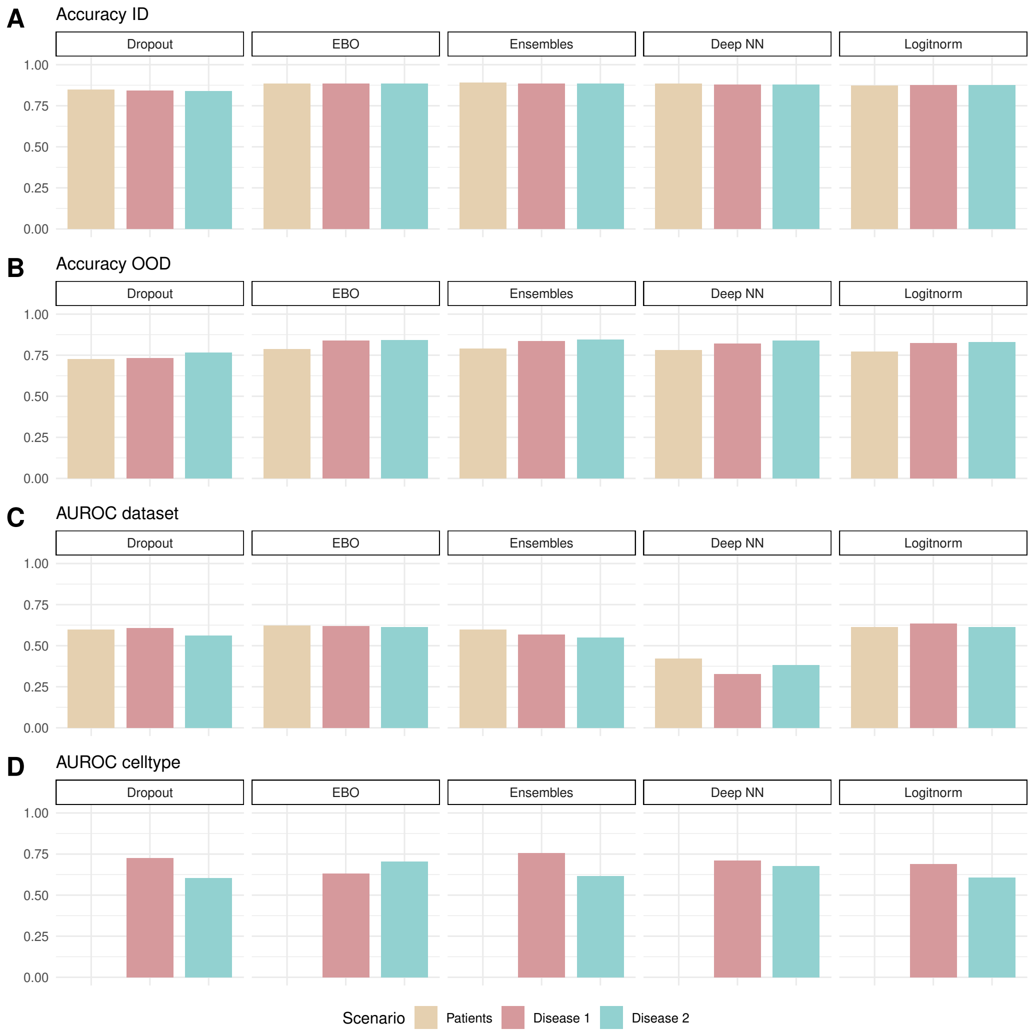


**Figure A6.** Visualisation of the ID annotation and OOD annotation results of the COPD dataset, for every method across the data splits (for patient splits the overall average is reported).

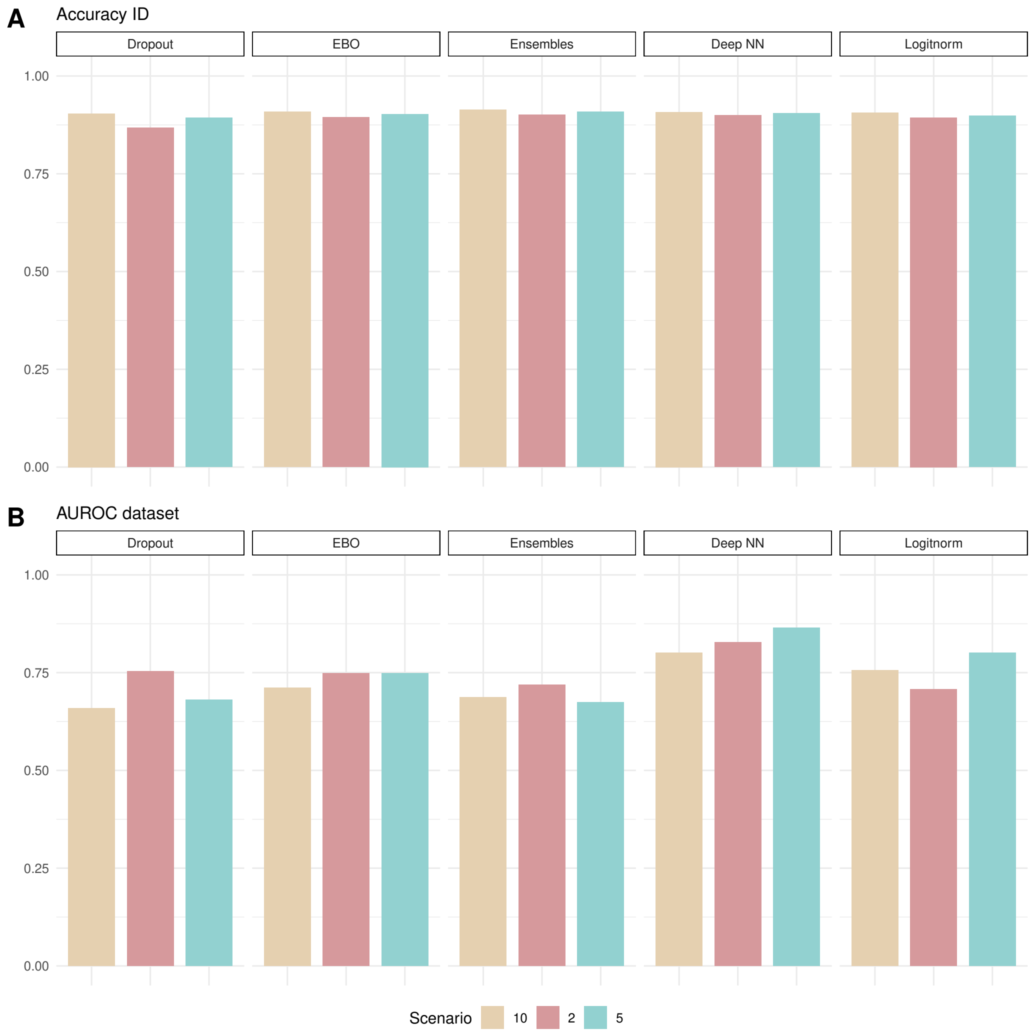


**Figure A7.** Visualisation of the ID annotation and OOD annotation results of the minor novel cell type prediction task, for every method across the data splits.
